## Supplementary figures and images for "An updated end-to-end ecosystem model of the Northern California Current reflecting ecosystem changes due to recent marine heat waves"

### Fig S1

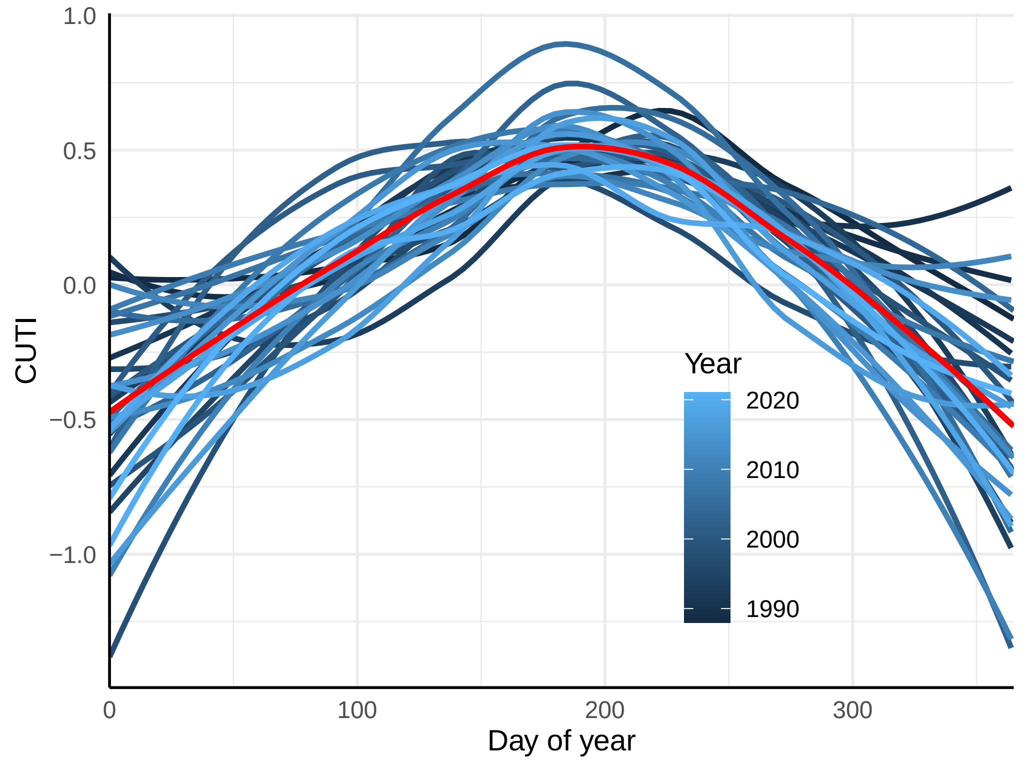
